# Cell Dynamics in Tumor Environment After Treatments

**DOI:** 10.1101/080895

**Authors:** Leili Shahriyari

**Affiliations:** Mathematical Biosciences Institute, The Ohio State University

## Abstract

Although the failure of cancers treatments has been mostly linked with the existence of resistant cells or cancer stem cells, new findings show a significant correlation between circulating inflammatory biomarkers and treatment failures. Most cancer treatments cause necrotic cell deaths in the tumor microenvironment. Necrotic cells send signals to the immune cells to start the wound healing process in the tissue. Therefore, we assume after stopping treatments there is a wound that needs to be healed. The stochastic simulations of epithelial cell dynamics after a treatment, which only kills cells without changing the tumor’s inflammatory environment, show that higher fitness of cancer cells causes earlier relapses. Moreover, the tumor returns even if a single cancer cell with high fitness remains in the wound’s boundary after such treatments. Although the involvement of cancer cells in the wound healing after treatments lead to the fast relapse, the cancer cells outside of the wound can also cause a slow recurrence of the tumor. Therefore, the absence of relapse after such treatments implies a slow-developing tumor that might not reach an observable size in the patients’ life time. Conversely, a large solid tumor in a young patient suggests the presence of high fitness cancer cells and therefore a high likelihood of relapse after conventional therapies. Additionally, the location of remaining cancer cells after treatments is a very important factor in the recurrence time. The fastest recurrence happens when a high fitness cancer cell is located in the middle of the wound. However, the longest time to recurrence corresponds to cancer cells located outside of the wound’s boundary.

## Introduction

Failure of traditional cancer therapies has been observed in almost all inflammatory cancers, and the high level of circulating inflammatory biomarkers is highly associated with the failure of treatments [1]. Some scientists argue that the reason behind the treatments’ failures might be the existence of resistant cells or cancer stem cells [2]. Here, we suggest another possible reason, which is based on the wound healing process in the tumor micro-environment after the treatments. Unnaturally dying cells send signals to the immune system to replace them and cure the wound. One of these damage-associated molecular pattern (DAMP) molecules, which triggers inflammation and immunity, is extracellular high mobility group box 1 (HMGB1) [3]. HMGB1 is passively released from necrotic cells, or actively secreted from stressed cancer cells and immune cells [4]. It has been observed that the release of HMGB1 in response to chemotherapy in leukemia increases resistance to the therapies [5]. Moreover, binding of HMGB1 to toll-like receptor 4 (TLR4) on dendritic cells (DCs) causes early recurrence after chemotherapy in breast cancer patients [6]. High levels of HMGB1 have also been observed in patients with non-small cell lung cancer (NSCLC) after tumors removed by surgery. In addition, significantly high levels of both HMGB1 and transition factor p65 were seen in NSCLC tumors with node metastasis [7]. Nasopharyngeal carcinoma (NPC) patients with high levels of HMGB1 expression had poor overall, disease-free survival [8]. These insights imply that most common cancer therapies such as surgery, radiation, and chemotherapy cause necrotic cell death [9], which activates the immune system in the same way as the wound-healing process [10].

In addition, in one experiment, human ovarian cancer cells were added to bone marrow cells recovered from irradiated mice with 1000 cGy. The irradiated bone marrow cells significantly increases proliferation of human ovarian cancer cells compared to non-irradiated ones [11]. Furthermore, micro-metastases in bone marrow are frequently observed after chemotherapy, and their existence significantly reduces the survival rate [12].

In summary, the most common cancer therapies generate a wound in the tumor by producing necrotic cell death. Necrotic cells in the tumor microenvironment activate the immune system to initiate the wound healing process. In many tumors, epithelial cells are adapted to divide in a much higher rate than normal cells; for example, tumor suppressor genes are inactivated. Moreover, in some cancers, like colitis associated cancer, there are some immune deficiencies, and immune cells are adapted to send a high level of proliferation or angiogenesis signals [13,14]. If the tumor includes adapted tissue or/and adapted immune cells, these adapted cells start the wound healing process in the tumor microenvironment. Adapted activated immune cells send more signals of proliferation and/or angiogenesis than normal cells [15]. Furthermore, if there were adapted tissue cells, they would divide at a much higher rate in response to these signals than normal cells. Thus, not only would the tumor come back after the treatment, but it would also grow more aggressively.

Recently, several mathematical models have been designed to study cell dynamics in normal tissue as well as tumors. Some stochastic models were also developed to investigate the cell dynamics in the process of tumor formation [16–23]. Although there are mathematical models studying the wound healing process [24–26], there are not many computational studies about the wound healing process after stopping treatments. In this paper, we develop two stochastic models (spatial and non-spatial) for cell dynamics after a treatment, which kills epithelial cells. We apply a stochastic model, because cell dynamics are stochastic, but we also obtain a deterministic model which approximately predicts the results of the non-spatial stochastic algorithm. We assume after the treatment there is a wound that needs to be healed. We denote the fitness of cancer cells over the normal cells by *r*. Briefly, assuming two cells: one cancerous and one normal, receive proliferation signals to fill out an empty location, then the probability that the cancer cell divides is *r* times the normal one. Vermeulen et al. [27] obtained the probability *P*_*R*_ that a mutant stem cell replaces its neighbor for various mutants; *P*_*R*_(*Kras*^*G*12*D*^ *v.s. WT*) = 0.78, *P*_*R*_(*Apc*^*+/*−^ *v.s. WT*) = 0.62, *P*_*R*_(*Apc*^−/−^ *v.s. WT*) = 0.79, under the normal condition *P*53^*R*172*H*^ did not confer a benefit 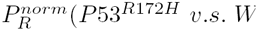 = 0.48, however in colitis 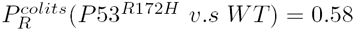. The fitness *r* in our model is given by 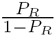, thus *r*(*Kras*^*G*12*D*^) = 3.5, *r*(*Apc*^+/−^) = 1.6, *r*(*Apc*^−/−^) = 3.8, *r*^*norm*^(*P*53^*R*172*H*^) = 0.9, and *r*^*colits*^(*P*53^R172H^) = 1.4. Therefore, in this work the fitness of cancer cells are assumed to be *r* = 3.8 (advantageous), *r* = 1 (neutral), and *r* = 0.9 (disadvantageous).

## Materials and Methods

After stopping a cancer treatment, which killed many cells, there is a wound that needs to be healed. In the wound healing process, necrotic cells as well as immune cells send signal to the nearby cells to divide and repair the wound. Moreover, some nearby epithelial cells are migrated to the wound with the help of platelets. Platelets also send some proliferation signals to these migrated cells [28].

Two stochastic models (non-spatial and spatial) are developed to simulate the recovery of cells after a treatment, which kills most of the cancer cells. The number of cancer cells and non-cancer cells at a given time t are respectively denoted by *C*(*t*) and *N*(*t*). The model’s assumption is that after treatments there is a wound, and cells start to divide to heal the wound. In other words, we assume the tissue needs approximately *D* number of cells, and the wound healing stops when the tissue reaches its desired number *D*. At the initial time of simulations, which is right after stopping treatments, there is a wound. Therefore, the initial number of cells, which is *C*(0) + *N*(0), is less than *D*. Thus, to heal the wound *D — C*(0) — *D*(0) cells are needed to be produced. At each updating time step, with probability 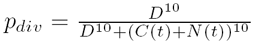 a cell divides, and with probability 1 — *p*_*div*_ a cell dies. The probability division function *p*_*div*_ is designed in such a way that if the total number of cells is less than *D*, then the division rate is much higher than death rate. However, when the total number of cells is approximately *D*, then the probability that a division happens is the same as the probability that a death occurs. Moreover, when a cell dies, with probability 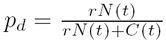, a non-cancer cell dies, or a cancer cell dies with probability 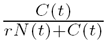. That means, higher fitness (i.e. higher probability of division) of cancer cells leads to the lower probability of their death. In this simulation, each updating step is the time that a change happens in the whole system, i.e. a cell divides or a cell dies. If a cell cycle is approximately 2 days, then the updating time *t* corresponds to approximately 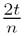days, where *n* is the total number of cells.

### Non-spatial model

The ratio of fitness of cancer cells to the normal cells is denoted by *r*. That means if a division happens at updating time *t*, with probability 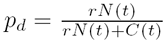, a cancer cell divides and with probability 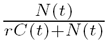, a non-cancer cell divides. At each updating time step, we run the following algorithm:

- With probability 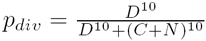a cell divides:
  - With probability 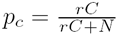, this division is the division of a cancer cell.
  - Or, with probability 1 — *p*_*c*_, this division is the division of a non-cancer cell.
- Or, with probability 1 — *p*_*div*_ a cell dies:
  - With probability 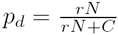, this death is the death of a non-caner cell.
  - Or, with probability 1 — *p*_*d*_, it is the death of a cancer cell.

After repeating the above algorithm for *T* updating time steps, we calculate the ratio of number of mutants over the total number of cells. Since this simulation is a stochastic model, we run the whole algorithm 10,000 times and we obtain the mean and standard deviations.

At each updating time step *t*, the number of cancer cells *C*(*t*) and non-cancer cells *N*(*t*) can be approximately obtained by the following deterministic system of equations.

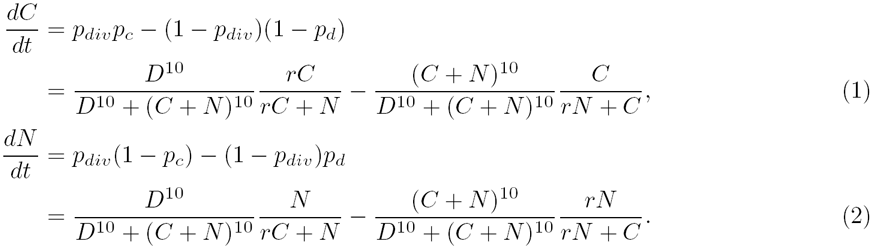

At the steady state of this system of equations, all cells are cancer cells (i.e. *N* = 0 and *C* = *D*), if *r* > 1, and if *r* < 1, then all cells become normal cells, (i.e. *N* = *D* and *C* = 0).

### Spatial model

A two dimensional lattice for the tissue is designed. The assumption is cells at the middle of the lattice are missing because of treatments. Note, necrotic cells send signals to the immune cells to start the wound healing process. Moreover, necrotic cells directly send signals of proliferations to the nearby cells. These proliferation signals diffuse over the neighborhood of the necrotic cells. For this reason, in this algorithm, only cells in the neighborhood of the empty spaces are dividing to replace missing cells. In other words, if there is an empty location, then any available cell located in the radius *ρ* from this empty space has a chance to divide. For example, if *ρ* =1, and the cell at the location (*i, j*) is missing, then any available cell at the locations {(*i* — 1, *j* — 1), (*i* — 1, *j*), (*i* — 1, *j* + 1), (*i, j* — 1), (*i, j* + 1), (*i* + 1, *j* — 1), (*i* + 1, *j*), (*i* + 1, *j* + 1)} has a chance to divide to fill out the location (*i, j*).For instance in Figure 1 (a), if *ρ* = 1, only cells located at {(3, 3)} ⋃ {(*i*, 2): 3 **≤** *i* **≤** 17} ⋃ {(*i*, 17): 2 **≤** *i* **≤** 17**}** ⋃ {(2,*j*): 2 **≤** *j* **≤** 17} ⋃ {(17, *j*): 2 **≤** *j* **≤** 17} have a chance to divide in the first updating time step. For simplicity, we assume the neighborhood size is fixed in the entire time of simulations. In other words *ρ* stays constant during the wound healing process and after the wound has been healed. In Figure 1 (a), the neighborhood of radius *ρ* = 1 and *ρ* = 3 of an empty space has been shown.

**Figure 1.**
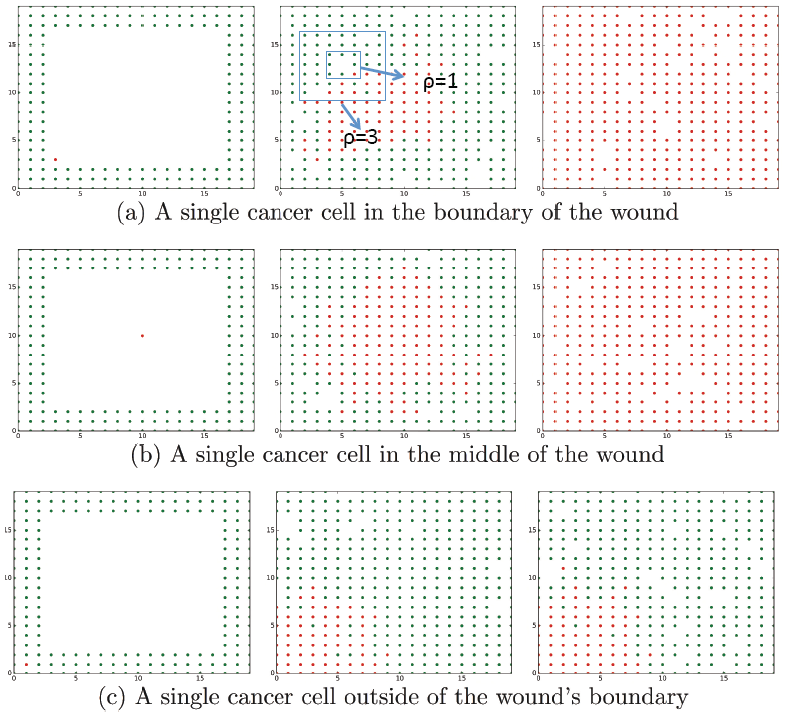
Spatial model. This figure shows cell dynamics in the wound healing process after treatments. At the initial time of simulations, a single cancer cell is located in the boundary of the wound (a), in the center of the wound (b), or outside of the wound’s border (c). Left plots show the initial time of simulations. Plots in the middle and right respectively show the healing time and the final time of simulations, which is *T* = 2000 number of updating time steps. Red and green circles are cancer cells and normal cells, respectively. The other parameters’ values are *r* = 3.8, *ρ* =1 and *μ* = *q* = 0. In (a), the neighborhood of radius *ρ* =1 and *ρ* = 3 of an empty space has been shown.

Here, we assume some percentage (%*q*) of cells are quiescent, that means they neither divide nor die. Therefore, we have two types of cells active and quiescent cells in the spatial model. Note that we treat both quiescent cancer and normal cells the same. In fact, we can assume there is no cancer quiescent cells, because they do not migrate and their number will stay the same. Hence, cancer quiescent cells would not change the ratio of cancer cells. For the similar reason, we do not consider any quiescent cells in the non-spatial model, because in the non-spatial model, cells only divide and die and there is no concept of neighborhood or migration. We also include possibility of cell migration from outside of the wound to the wound during the wound healing process. Because epithelial cells are mostly static, and epithelial-mesenchymal transition (EMT) occurs during some particular situations such as wound healing process or tumor invasion [29–31], we assume cells stop migrating after the wound has been healed. In other words, at each updating time step till the time that the tissue reaches its desired number of cells, with probability *μ* an active cell migrates, or with probability (1 — *μ*) a cell division or death occurs. Thus, the algorithm for the spatial model is the following: we set the *Heal* = *False* at the initial time, then at each updating time step *t*, we run the following sub-algorithm:

- if *Not*(*Heal*), then with probability *μ* a uniformly randomly chosen active cell migrates to a uniformly chosen active location randomly chosen empty location.
- if *Heal* or (if *Not*(*Heal*), with probability 1 — *μ*), then one of the following scenarios happens:
  - With probability 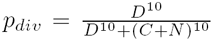, an empty location, which has at least one active cell in its neighborhood of radius *ρ*, is randomly uniformly chosen. Then, a cell located in the neighborhood of the empty location divides:
    - With probability 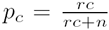, this division is the division of a cancer cell, where c and n are respectively the number of cancer and active normal cells in the neighborhood of the empty location.
    - Or, with probability 1 — *p*_*c*_, this division is the division of an active non-cancer cell located in the neighborhood of radius ρ from the empty location.
  - Or, with probability 1 — *p*_*div*_ a cell dies:
    - With probability 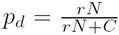, this death is the death of a non-caner cell, where N and C are respectively the total number of active normal and cancer cells in time *t*.
    - Or, with probability 1 — *p*_*d*_, it is the death of a cancer cell.
- if *C*(*t*) + *N*(*t*) = *D: Heal* = *True*

In this model, the only interaction between cancer cells and normal cells is their competition to fill out available empty spaces and their probability of death. For example, if there are n number of active normal cells and m number of cancer cells in the neighborhood of an empty space, then with probability 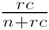, one cancer cell will divide; and with probability 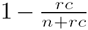, one normal cell will win the competition to divide and fill out the empty space. Additionally, if there are *N* number of active normal cells and *C* number of cancer cells in the entire system, then with probability 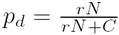 an active normal cell dies, and with probability 1 — *p*_*d*_ one cancer cell dies.

Although values of models’ parameters might vary for different tumors and treatments, they might have some specific range. However, because of the lack of biological data, in simulations, we choose a wide range of values for parameters. Here, the best-case scenario that can happen after the treatments is modeled in both non-spatial and spatial simulation. In the best-case scenario, after the wound has been healed cells follow the normal homeostasis, which is going toward a relative stable equilibrium. In other words, the number of cells stays approximately constant, i.e. the death rate is approximately the same as division rate. Note the probability of division is given by the function *p*_*div*_. This function is defined so that when the total number of cells reaches the desired number of cells (*D* = *C*(*t*) + *N*(*t*)), then the probability of division becomes the same as the probability of death (= 0.5).

## Results

In this work, we model the cell dynamics in the tumor microenvironment after stoping treatments, which cause a wound in the tissue. Right after stopping treatments, the division rate is high because the tissue wants to reach its normal concentration. Although cells might not follow the normal cell division and death rates after the wound has been healed, but for simplicity, we assume the best-case scenario happens, i.e. going toward the normal homeostasis, which is having a relatively stable constant number of cells. In other words, the tissue follows the normal cell division and death rates after the wound has been healed. Therefore, we denote the time that more than 99% of the tissue cells are cancer cells as the recurrence time. In the spatial model, the wound is placed in the center of a grid and cells are located at the boundary of the wound. In order to obtain the effect of the location of cancer cells, at each simulation a single cancer cell is located in a specific location; center, border, or outside of the wound (See Figure 1). In this model, only cells located near an empty space are able to divide, and any cell can go toward the normal cell death. However, in the non-spatial model, all cells are able to divide and die. In both models, the number of empty spaces and the number of desired cells, i.e. the normal tissue’s size, is respectively denoted by *E* and *D*.

### High fitness of cancer cells leads to fast relapse

Our stochastic models show that the time to recurrence, which is the time that more than 99% of the tissue cells are cancer cells, is a decreasing function of the fitness of cancer cells. In other words, if cancer cells have high fitness then the tumor would reappear after stopping the treatments in a very short time. Figure 2(a) shows the effect of the fitness of cancer cells left after treatments in the time to relapse for non-spatial model. The disadvantageous cancer cells, i.e. the cancer cells with fitness less than one (the relative fitness of w.t. cells), will be removed from the tissue in the non-spatial model. However, the advantageous cancer cells, which have fitness more than one, will always take over the wound and then the entire tissue in the non-spatial model. High fitness cells that are involved in the wound healing process will rapidly divide, and fill out the empty spaces. Moreover, in the spatial model, if an advantageous cancer cell is located close to an empty space generated from a normal cell death, then the progeny of the cancer cell will fill out the empty space with a high probability (See Figure 1(c)). Note, necrotic cells send proliferation signals to the nearby cells, and high fitness cells are more likely to respond to these signals and divide (See Figure 1).

**Figure 2.**
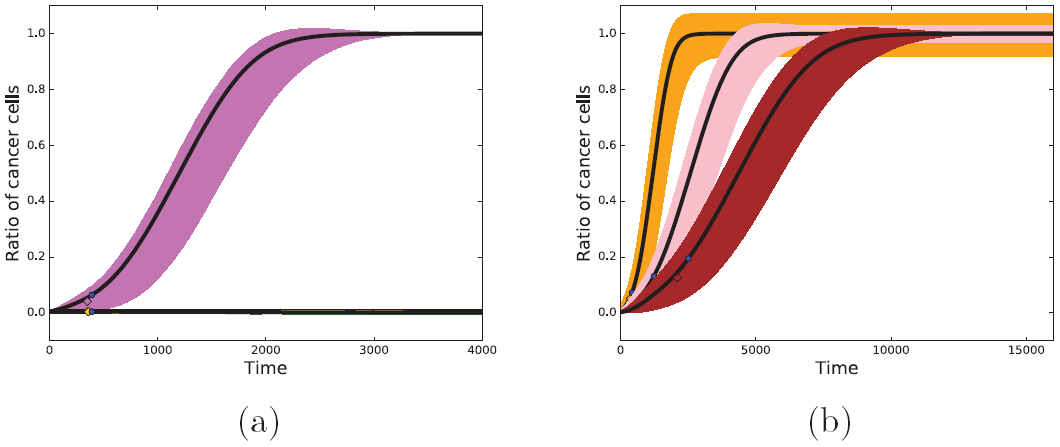
Non-spatial model. This figure presents the ratio of cancer cells over the total number of cells as a function of time, i.e. the *x*—axis is the number of updating steps. In this plot, the blue circles (the result of the formula) and diamonds (the result of simulations) indicate the healing time, i.e. the first time *t* that total number of cells reaches the desired number, *N*(*t*) + *C*(*t*) = *D.* The black lines show the solutions of the system of equations. In sub-figure (a), at the initial time there are 195 normal cells and a single cancer cell, and the desired number of cells is approximately *D* = 400. The gold, green, and orchid colors show the average and standard deviation of 100 independent runs for *r* = 0.9, *r* = 1.0, and *r* = 3.8, respectively. The ratio of cancer cells is around zero for *r* = 0.9 and *r* = 1.0. In sub-figure (b), the fitness of cancer cells is *r* = 3.8. The orange, pink, and brown colors respectively show the mean plus/minus standard deviation of 1000 independent runs for (*N*(0), *D*) = (80,400), (575, 900), and (1155, 1600). Where *N*(0) is the initial number of normal cells, and *D* is the desired number of cells.

Figure 3 shows the results of 100 independent individual runs for the advantageous (*r* > 1), neutral (*r* = 1), and disadvantageous (*r* < 1) cancer cells. Figure 3(a) indicates that the wound heals faster if the cancer cells are advantageous and they are involved in the wound healing process. Moreover, if advantageous cancer cells are involved in the healing process, then their progeny will rapidly take over the entire tissue. However, the disadvantageous cancer cells located in the border of the wound or outside of the wound’s boundary are not able to colonize, and will be eventually washed out from the tissue.

### Small wounds and fast recurrence

In order to simulate the effect of the wound’s size in the recurrence time, we change the grid’s size in the spatial model in Figures 4 and 5. In other words, we create a large grid for a large wound, and we consider three layers of cells around the wound in Figures 4 and 5 (See Figure 1). To compare the spatial and non-spatial model in Figure 5, we consider the same number of cells and empty spaces for the non-spatial model at the initial time of simulations as the spatial model. Expectedly, simulations show that small wounds heal fast (See Figure 4). However, the ratio of cancer cells over the total number of cells after healing depends on the location and the fitness of cancer cells as well as the wound’s size. The non-spatial model shows that time to recurrence is an increasing function of the wound size (See Figure 2(b)). However, the spatial model indicates that if cancer cells are located at the wound’s boundary or outside of the wound, then the smaller wounds relapse faster than larger ones if the probability of cell migration is small (See Figures 5 and 6). Note, when the wound is small, i.e. a small number of cells are needed, then the wound heals fast and then the tissue follows the normal cell death and division rates quickly. Then, cancer cells outside of the wound or in the wound’s boundary, which were not involved in the healing process, gets a chance to divide and take over the tissue. Thus, cancer cells, which were not involved in the wound healing process, in smaller wounds gets a chance to divide faster than in a larger wound, because a larger wound needs more time to heal. Note that the time to recurrence is more sensitive to the fitness of cancer cells than the location of cancer cells and the size of the wound (See Figure 5). Moreover, cell migration during the wound healing process increases the chance of involvement of cancer cells, which are not located in the middle of the wound (See Figure 6). Figure 6 also shows that the maximum ratio of cancer cells at the final time of simulations corresponds to the maximum of the migration probability and wound’s size. In other words, if migration probability is high and cancer cells are outside of the wound, then larger wounds leads to a faster relapse than smaller tumors. Thus, when we see earlier relapses after removing large tumors (leading to large wounds) more often than small ones, it is because cancer cells in larger tumors have higher fitness than smaller ones or there is a higher chance that cancer cells are involved in the wound healing after treating larger tumors.

**Figure 3.**
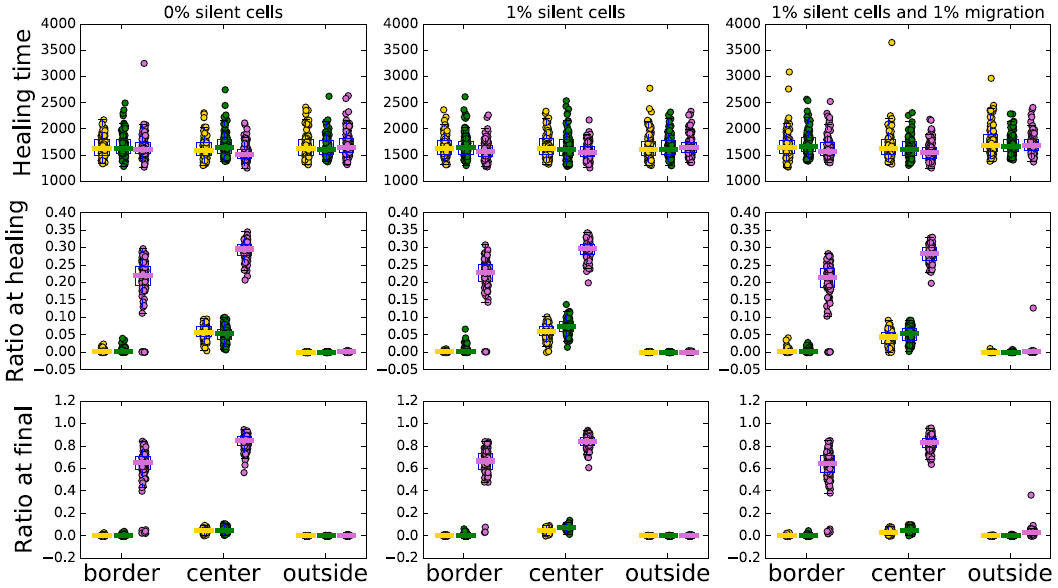
Cell dynamics in the spatial model. The sub-figures located at the top of this figure show the healing time, while the sub-figures in the center and bottom indicate the ratio of cancer cells over the total number of cells at the healing time and final time, respectively. In each of these sub-figures, the result of 100 individual runs is shown by circles when a single cancer cell is located in the border, center, and outside of the wound at the initial time of simulations. The orchid, green, and gold colors correspond to the fitness of cancer cells *r* = 3.8, *r* = 1.0, and *r* = 0.9, respectively. In this figure, *E* = 1156, *D* = 1600, and *ρ* = 1 The other parameters’ values are *μ* = *q* = 0 for plots in the left column, *q* = %1 and *μ* = 0 for the plots in the middle, and *q* = %1 and *μ* = 0.01 for the plots in the right.

### Location of cancer cells is important

In the spatial simulations, a single cancer cell is placed in different locations. If an advantageous cancer cell is involved in the wound healing process, i.e. it is located in the middle of the wound or at the wound’s border, then it would outcompete normal cells in the wound healing process. Therefore, the tumor reoccurs very quickly, because after the wound has been healed there are many cancer cells that are involved in the tissue’s normal homeostasis (See Figures 5 and 6). However, if the advantageous cancer cell is located outside of the wound and they are not involved in the wound healing process, then the wound is healed by normal cells. In this case, the tumor would slowly re-generate because of the cancer cell’s involvement in the normal cell division in the tissue after the wound has been healed (See Figures 5 and 6).

**Figure 4.**
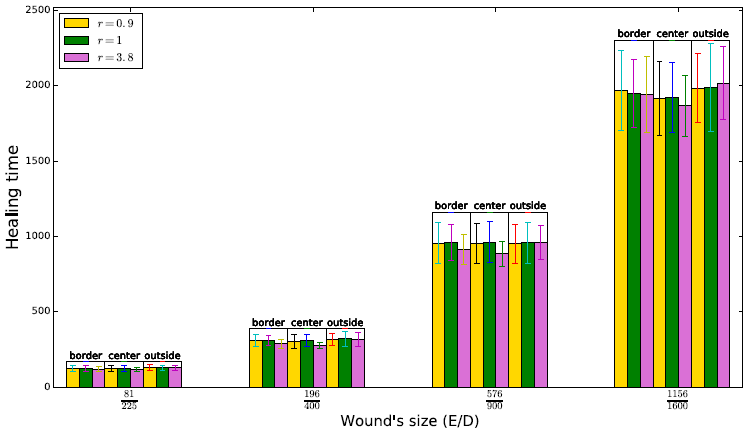
Healing time. This figure indicates the first time that the wound is healed, i.e. the total number of cells is approximately the same as the desired number of cells, when a single cancer cells is located at the border, center, and outside of the wound in the initial time of simulations. In this figure, the orchid, green, and gold colors correspond to the fitness of cancer cells *r* = 3.8, *r* = 1.0, and *r* = 0.9, respectively. The other parameters’ values are *ρ* =1 and *μ* = *q* = 0.

**Figure 5.**
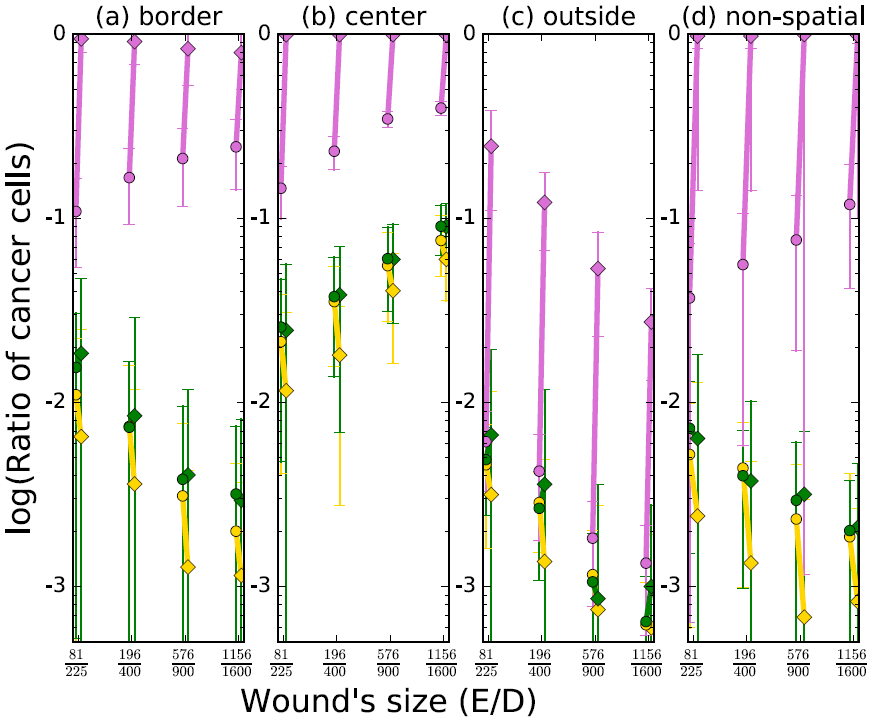
Ratio of cancer cells. This figure shows the ratio of the number of cancer cells over the total number of cells. The circles and diamonds indicate the ratio of cancer cells at the healing time and the final time of simulations, respectively. At the initial time of these simulations, a single cancer cell is located in the boundary of the wound (sub-figure (a)), in the center of the wound (sub-figure (b)), or outside of the boundary of the wound (sub-figure (c)). The final time of simulations, which corresponds to *T* = 10*D* number of updating time steps, where *D* is the tissue’s size, i.e. the desired number of cells. In this figure, the *x*-axis is the ratio of *E/D,* where *E* is the number of empty locations. The orchid, green, and gold colors correspond to the fitness of cancer cells *r* = 3.8, *r* = 1.0, and *r* = 0.9, respectively. The other parameters’ values are *ρ* = 1 and *μ* = *q* = 0.

**Figure 6.**
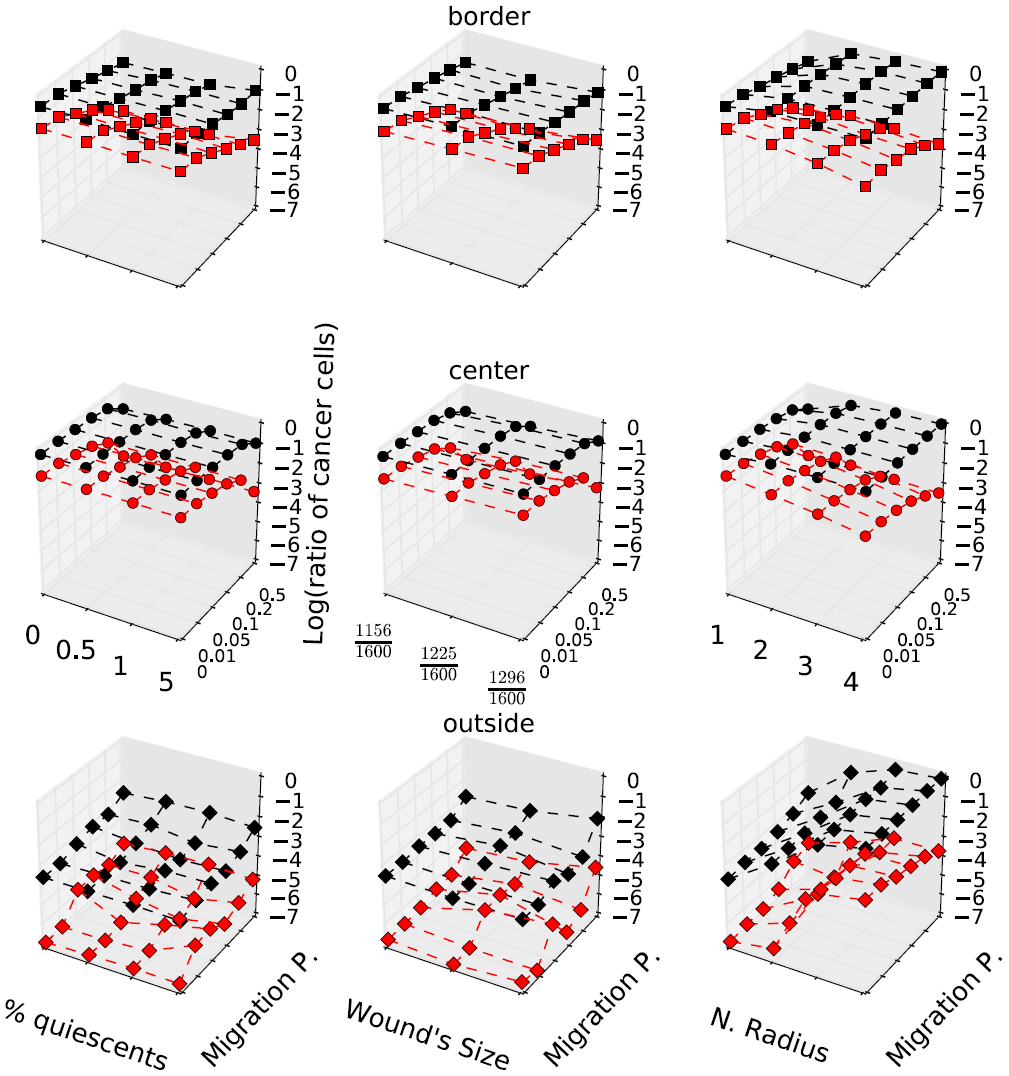
Ratio of cancer cells in spatial model. This figure shows the log of the mean of ratio of the number of cancer cells over the total number of cells for 100 independent runs. The red and black colors indicate the ratio of cancer cells at the healing time and the final time of simulations, respectively. At the initial time of these simulations, a single cancer cell is located in the boundary of the wound (top sub-figure), in the center of the wound (middle sub-figures), or outside of the boundary of the wound (bottom sub-figures). The final time of simulations, which corresponds to *T* = 10*D* number of updating time steps, where *D* is the tissue’s size, i.e. the desired number of cells. In this figure, the wound’s size is the ratio of *E/D,* where *E* is the number of empty locations. The parameters’ values for this figure are *r* = 3.8, and in left sub-figure, we vary the migration probability and the percentage of quiescent cells and other parameters are *ρ* =1 and 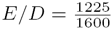. In the middle sub-figures, we vary the wound’s size and migration probability, and other parameters are *ρ* =1 and *q* = %1. In the right sub-figures, we vary the neighborhood radius and migration probability, and other parameters are *q* = %1, and 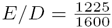, if they were not varied in the plots.

### The tissue’s size is as important as the wound’s size

After healing, the percent of cancer cells involved in the normal homeostasis is a very important factor in the recurrence time. If the wound is large and cancer cells are involved in the wound’s healing process, then cancer cells will cover a high percent of the tissue. However, if cancer cells are involved in the healing of a very small wound, then cancer cells only cover a small portion of the tissue after healing. Moreover, if the cancer cells are not involved in the wound healing process, i.e. are located outside of the wound’s boundary and the probability of cell migration is small, then they are only involved in the normal homeostasis. A high percent of the cancer cells involved in the normal homeostasis leads to a fast recurrence (See Figures 5 and 6). For this reason, if a single cancer cell is located outside of the wound’s border and probability of cell migration is small, then the tumor reoccurs faster when the tissue’s size is smaller. Furthermore, when the tissue’s size is fixed and advantageous cancer cells are not involved in the wound healing, then smaller wound leads to faster recurrence. Note, smaller wounds heal faster and as soon as the wound is healed, then tissue goes toward the normal homeostasis. Although a larger wound’s size delays the tumor recurrence, if tumor cells are located outside or in the boundary, the cell migrations would increase the ratio of cancer cells in the healing time for cancer cells located outside of a large wound (See middle sub-figures of Figure 6).

### Migrations during wound healing process and and the size of neighborhoods are very important in tumors recurrences

Cell migration during wound healing process increases the chance of involvement of cells located outside of the wound in the wound healing process. Therefore, if a cancer cell is located in outside of the wound, then the ratio of cancer cells increases when a high percent of cells are migrating during the wound healing process (See bottom sub-figures of Figure 6). However, cell migration increases the number of normal cells involving in the healing process, if there are no cancer cells outside of the wound. Thus, cell migrations decreases the chance of the tumor recurrence if all tumor cells are located inside or at the border of the wound (See top and middle sub-figures of Figure 6). For the same reasons, the neighborhood size have the same effect as the cell migrations; it increases the chance of involvement of cells, which are located outside of the wound (See right sub-figures of Figure 6). Furthermore, comparing the plots, which are located in the same columns in Figure 6, reveal that although migrations increase the ratio of cancer cells, however still cancer cells located at the middle and the boundary of the wound lead to a higher ratio of cancer cells at the final time of simulations. Only when both migration probability and neighborhood radius are very high (*μ* ≥ 0.5 and *ρ* ≥ 4), the ratio of cancer cells at final time starting from a single cancer cell located outside of the wound can become close to the ratio of the cancer cells starting from a single caner cell located inside the wound (See right sub-figures of Figure 6).

### The existence of quiescent cell increases the variation of the results

Although the different percentage of quiescent cells does not vary the mean of the ratio of cancer cells much, it leads to a higher standard variation of the ratio of cancer cells (See left sub-figures of Figure 6). The reason for observing this phenomena is that the locations of quiescent cells are chosen randomly. If most of the quiescent cells are located close to the wound’s boundary or in the neighborhood of the cancer cell, then the probability that the cancer cell divides increases. The ratio of cancer cells increases if there are many quiescent in the neighborhood of cancer cells, because quiescent cells do not compete with cancer cells to divide and filling out empty spaces. This phenomenon can be seen more clearly when cancer cells are located outside of the wound, because in this case the chance of quiescent cells being located in the neighborhood of the cancer cell increases (See left sub-figures of Figure 6). Quiescent cells cannot be in the neighborhood of the single cancer cell in the middle of the wound, because the neighborhood of the cancer cell located in the middle of the wound is empty places at the initial time and quiescent cells only appear in the initial time.

## Discussion

Since the disadvantageous cancer cells are not able to colonize, the development of the tumor before the treatment indicates that the fitness of cancer cells must be more than one. The stochastic simulations of cell dynamics show that if any cancer cells with fitness more than one remain after the treatments that only kill epithelial cells, then the tumor will relapse. However, no relapses have been detected after the treatment for some non-inflammatory cancers. It is very unlikely for any treatment to kill every single cancer cell. Therefore, if the cancer does not come back (treatments are working), then the fitness of cancer cells after the treatment should be slightly more than one, which is the relative fitness of w.t. cells. If the fitness of cancer cells is slightly more than one, then tumor will very slowly develops over time. This implies, if a treatment is working, then most likely the tumor is old. We can also conclude that if a young patient has a large solid tumor, i.e. the tumor developed in a short time, then tumor cells must have high fitness. Simulations also reveal that high fitness cancer cells correspond to the poor outcome, i.e. fast recurrences, which is consistent with clinical observations [32]. For these patients, common treatments, which kill all cells, can not increase the survival time significantly, because as soon as the treatment stops, tumor cells begin the healing process, get to the normal concentration, and grow.

Cell migrations and the diffusion of proliferation signals to a large neighborhood during the wound healing process increases the chance of involvement of cancer cells located outside of the wound. However, still the longest time to recurrence corresponds to cancer cells located outside of the wound’s boundary (See Figures 3 and 6). Furthermore, since there are more cells in the neighborhood of cancer cells located outside of the wound at the initial time of simulations, than cancer cells, which are located inside or in the boundary of the wound, a high percentage of quiescent cells increases the chance of having quiescent cells in the neighborhood of cancer cells located outside of the wound. Quiescent cells do not compete with cancer cells to fill out empty spaces, therefore the existence of quiescent cells would increase the chance of cancer cells’ divisions, if they are located in the neighborhood of cancer cells. Thus, existence of quiescent cells would increase the chance of division of cancer cells, which are located outside of the wound.

There are many reports about the occurrence of the metastasis to the surgical wounds [33, 34]. The incidence of wound metastasis provides evidence for the importance of the location of the tumor cells after the treatments. In agreement with the clinical observations, the stochastic simulations show that the involvement of cancer cells in the wound healing process after the treatments is a very important factor in the recurrence time. The simulations indicate that if the advantageous tumor cells are located in the middle of the wound, then the tumor will rapidly relapse. According to Vermeulen et al. [27], the fitness of mutant *P*53^*R*172*H*^ stem cells in normal conditions is not more than one, but it is more than one in inflammatory environments. Note, proliferation signals from immune cells and necrotic cells are a very important factor in the tumor growth. In this work, it is assumed that cells follow the normal homeostasis after the wound has been healed. However, in some cancers, the tissue does not follow the normal homeostasis after the wound has been healed, or the wound will be never healed. For example, if there is still an inflammation after filling out empty spaces in the tissue, then the level of inflammatory signals like proliferation signals can be very high in the tissue. That can lead to a high number of divisions of epithelial cells, causing re-growth of the tumor. Based on these insights, one can conclude that inflammation plays a more important role in the progression of tumors than mutations in epithelial cells. The inflammation not only can cause mutation in epithelial cells [14], but can also change their fitness. Thus, the effective treatment for inflammatory carcinomas must change the inflammatory environment of the tumor.

The developed model has some limitations such as assuming normal homeostasis after the tissue reaches its normal desired number of cells. Moreover, for simplicity, we have assumed that only a single cancer cell has been remained after treatments. However, this model can be generalized by assuming the existence of several cancer cells in different locations. Also, for simplicity we have assumed that the quiescent cells stay quiescent and all active cells stay active. For this reason, the model cannot include a high percentage of quiescent cells at the initial time of simulations. If the percentage of quiescent cells was higher than %5, then all neighbors of some empty places become quiescent cells. Then, that empty place could not be filled. Therefore, in simulations, we have assumed that the percentage of quiescent cells is less than %5. Therefore, the next step would be improving the model to overcome these limitations.

## References

1. Pierce BL, Ballard-Barbash R, Bernstein L, Baumgartner RN, Neuhouser ML, Wener MH, et al. Elevated Biomarkers of Inflammation Are Associated With Reduced Survival Among Breast Cancer Patients. Journal of Clinical Oncology. 2009;27(21):3437–3444.

2. Dhawan A, Kohandel M, Hill R, Sivaloganathan S. Tumour control probability in cancer stem cells hypothesis. PLoS ONE. 2014;9(5).

3. Lotze MT, Tracey KJ. High-mobility group box 1 protein (HMGB1): nuclear weapon in the immune arsenal. Nature reviews Immunology. 2005;5(4):331–342.

4. Wang H, Bloom O, Zhang M, Vishnubhakat JM, Ombrellino M, Che J, et al. HMG-1 as a late mediator of endotoxin lethality in mice. Science (New York, NY). 1999;285(5425):248–251.

5. Liu L, Yang M, Kang R, Wang Z, Zhao Y, Yu Y, et al. HMGB1-induced autophagy promotes chemotherapy resistance in leukemia cells. Leukemia: official journal of the Leukemia Society of America, Leukemia Research Fund, UK. 2011;25(1):23–31.

6. Apetoh L, Ghiringhelli F, Tesniere A, Criollo A, Ortiz C, Lidereau R, et al.. The interaction between HMGB1 and TLR4 dictates the outcome of anticancer chemotherapy and radiotherapy; 2007.

7. Zhang X, Wang H, Wang J. Expression of HMGB1 and NF-?B p65 and its significance in non-small cell lung cancer. Wspolczesna Onkologia. 2013;17(4):350–355.

8. Wu D, Ding Y, Wang S, Zhang Q, Liu L. Increased expression of high mobility group box 1 (HMGB1) is associated with progression and poor prognosis in human nasopharyngeal carcinoma. The Journal of pathology. 2008;216(2):167–175.

9. Zong WX, Thompson CB. Necrotic death as a cell fate; 2006.

10. Harless WW. Cancer treatments transform residual cancer cell phenotype. Cancer cell international. 2011;11(1):1.

11. Gunjal PM, Schneider G, Ismail AA, Kakar SS, Kucia M, Ratajczak MZ. Evidence for induction of a tumor metastasis-receptive microenvironment for ovarian cancer cells in bone marrow and other organs as an unwanted and underestimated side effect of chemotherapy/radiotherapy. Journal of Ovarian Research. 2015;8(1).

12. Braun S, Kentenich C, Janni W, Hepp F, de Waal J, Willgeroth F, et al. Lack of effect of adjuvant chemotherapy on the elimination of single dormant tumor cells in bone marrow of high-risk breast cancer patients. Journal of clinical oncology: official journal of the American Society of Clinical Oncology. 2000;18(1):80–86.

13. Terzic J, Grivennikov S, Karin E, Karin M. Inflammation and Colon Cancer. Gastroenterology. 2010;138(6):2101–2114.

14. Waldner MJ, Neurath MF. Colitis-associated cancer: The role of T cells in tumor development. Seminars in Immunopathology. 2009;31(2):249–256.

15. Shahriyari L. A new hypothesis: some metastases are the result of inflammatory processes by adapted cells, especially adapted immune cells at sites of inflammation. F1000Research. 2016 feb;5:175.

16. Enderling H, Chaplain MAJ, Anderson ARA, Vaidya JS. A mathematical model of breast cancer development, local treatment and recurrence. Journal of theoretical biology. 2007 may;246(2):245–59.

17. Komarova NL, Shahriyari L, Wodarz D. Complex role of space in the crossing of fitness valleys by asexual populations. Journal of The Royal Society Interface. 2014 mar;11(95):20140014–20140014.

18. Shahriyari L, Komarova NL. Symmetric vs. Asymmetric Stem Cell Divisions: An Adaptation against Cancer? PLoS ONE. 2013 oct;8(10):e76195.

19. Shahriyari L, Komarova NL. The role of the bi-compartmental stem cell niche in delaying cancer. Physical Biology. 2015 jul;12(5):055001.

20. McHale PT, Lander AD. The protective role of symmetric stem cell division on the accumulation of heritable damage. PLoS computational biology. 2014 aug;10(8):e1003802.

21. Jilkine A, Gutenkunst RN. Effect of dedifferentiation on time to mutation acquisition in stem cell-driven cancers. PLoS computational biology. 2014 mar;10(3):e1003481.

22. Shirayeh AM, Shahriyari L. New Insights into Initiation of Colon and Intestinal Cancer: The Significance of Central Stem Cells in the Crypt. arXiv preprint arXiv:161004089. 2016;.

23. Shahriyari L, Shirayeh AM. Optimal structure of heterogeneous stem cell niche: The importance of cell migration in delaying tumorigenesis. bioRxiv. 2016;p. 082982.

24. Maini PK, McElwain DLS, Leavesley DI. Traveling wave model to interpret a wound-healing cell migration assay for human peritoneal mesothelial cells. Tissue engineering. 2004;10(3-4):475–82.

25. Maggelakis SA. A mathematical model of tissue replacement during epidermal wound healing. Applied Mathematical Modelling. 2003;27(3):189–196.

26. Olsen L, Sherratt JA, Maini PK. A mathematical model for fibro-proliferative wound healing disorders. Bulletin of Mathematical Biology. 1996 jul;58(4):787–808.

27. Vermeulen L, Morrissey E, van der Heijden M, Nicholson AM, Sottoriva A, Buczacki S, et al. Defining Stem Cell Dynamics in Models of Intestinal Tumor Initiation. Science. 2013 nov;342(6161):995–998.

28. Nurden AT, Nurden P, Sanchez M, Andia I, Anitua E. Platelets and wound healing. Frontiers in bioscience: a journal and virtual library. 2008 may;13:3532–48.

29. Zvaifler NJ. Relevance of the stroma and epithelial-mesenchymal transition (EMT) for the rheumatic diseases. Arthritis research & therapy. 2006;8(3):210.

30. Lu M, Jolly MK, Onuchic J, Ben-Jacob E. Toward Decoding the Principles of Cancer Metastasis Circuits. Physics in Cancer Research Cancer Res. 2014;74(17):4574–87.

31. Jolly MK, Huang B, Lu M, Mani SA, Levine H, Ben-Jacob E. Towards elucidating the connection between epithelial-mesenchymal transitions and stemness. Journal of the Royal Society, Interface / the Royal Society. 2014;11(101):20140962.

32. Gabriel Ca, Domchek SM. Breast cancer in young women. Breast Cancer Research. 2010;12(5):212.

33. Curet MJ. Port site metastases. American Journal of Surgery. 2004;187(6):705–712.

34. David W, Mathew G, Ellis T, Baigrie CF, Rofe AM, Jamieson GG. Gasless Laparoscopy May Reduce the Risk of Port-Site Metastases Following Laparascopic Tumor Surgery. Archives of Surgery. 1997 feb;132(2):166.

